# Avoiding sedentary behaviors requires more cortical resources than avoiding physical activity: an EEG study

**DOI:** 10.1101/277988

**Authors:** Boris Cheval, Eda Tipura, Nicolas Burra, Jaromil Frossard, Julien Chanal, Dan Orsholits, Rémi Radel, Matthieu P. Boisgontier

## Abstract

- Individuals are slower at approaching sedentary than physical activity stimuli
- Individuals are quicker at avoiding sedentary than physical activity stimuli
- These effects are particularly pronounced in physically active individuals
- Avoiding sedentary behaviors is associated with high levels of conflict monitoring and inhibition
- Additional brain resources are required to escape a general attraction toward sedentary behaviors

**Abstract:** Why do individuals fail to exercise regularly despite knowledge of the risks associated with physical inactivity? Automatic processes regulating exercise behaviors may partly explain this paradox. Yet, these processes have only been investigated with behavioral paradigms based on reaction times. Here, using electroencephalography, we investigated the cortical activity underlying automatic approach and avoidance tendencies toward stimuli depicting physical activity and sedentary behaviors in 29 young adults who were physically active (n=14) or physically inactive but with the intention of becoming physically active (n=15). Behavioral results showed faster reactions when approaching physical activity compared to sedentary behaviors and when avoiding sedentary behaviors compared to physical activity. These faster reactions were more pronounced in physically active individuals and were associated with changes during sensory integration (earlier onset latency and larger positive deflection of the stimulus-locked lateralized readiness potentials) but not during motor preparation (no effect on the response-locked lateralized readiness potentials). Faster reactions when avoiding sedentary behaviors compared to physical activity were also associated with higher conflict monitoring (larger early and late N1 event-related potentials) and higher inhibition (larger N2 event-related potentials), irrespective of the usual level of physical activity. These results suggest that additional cortical resources were required to counteract an attraction to sedentary behaviors. Data and Materials [https://doi.org/10.5281/zenodo.1169140].

## 1. Introduction

Why do we fail to exercise regularly (Kohl, et al., 2012) despite the known negative effects of physical inactivity on health (e.g., Ekelund, et al., 2016; Lee, et al., 2012)? This exercise paradox could be explained by an imbalance between controlled and automatic processes, which have been defined in dual-process models of health behaviors (Brand & Ekkekakis, 2017; Hofmann, Friese, & Wiers, 2008). Controlled processes are initiated intentionally, require cognitive resources, and operate within conscious awareness. Automatic processes are initiated unintentionally, tax cognitive resources to a much lesser extent, occur outside conscious awareness, and can be problematic when they come into conflict with controlled processes (Marteau, Hollands, & Fletcher, 2012; Strack & Deutsch, 2004). For example, the detection of an opportunity for being sedentary can automatically trigger a drive competing with the conscious intention to adopt a physically active behavior, thereby disrupting or preventing its implementation. While the dichotomization proposed by dual-process models has been subject to debate (Melnikoff & Bargh, 2018), this pragmatic simplification has facilitated the integration of findings from heterogeneous concepts and experimental designs. An increasing number of studies show that automatic reactions, such as attentional capture (Berry, 2006; Berry, Spence, & Stolp, 2011; Calitri, Lowe, Eves, & Bennett, 2009), affective reactions (Antoniewicz & Brand, 2016; Bluemke, Brand, Schweizer, & Kahlert, 2010; Chevance, Caudroit, Romain, & Boiché, 2017; Conroy, Hyde, Doerksen, & Ribeiro, 2010; Rebar, Ram, & Conroy, 2015), and approach tendencies (Cheval, Sarrazin, Isoard-Gautheur, Radel, & Friese, 2015; Cheval, Sarrazin, Isoard-Gautheur, Radel, & Friese, 2016; Cheval, Sarrazin, & Pelletier, 2014) are important for the regulation of exercise behaviors (see Cheval, et al., 2018; Rebar, et al., 2016; Schinkoeth & Antoniewicz, 2017, for reviews). However, these studies have mainly focused on automatic reactions triggered by physical activity, whereas only few studies have examined automatic reactions triggered by sedentary behaviors or behaviors minimizing energetic cost.

We define energetic cost minimization as the automatic processes aiming to achieve the most cost-effective behavior. Energetic cost minimization is considered a fundamental principle in multiple fields such as exercise physiology and biomechanics but has been completely disregarded in the field of exercise psychology. Its impact on behavior is clearly illustrated in gait, where it determines the moment we switch from walking to running (Srinivasan & Ruina, 2006). In a recent systematic review, we contend that including the concept of energetic cost minimization into the dominant approaches to exercise behavior can improve our understanding of the exercise paradox (Cheval, et al., 2018). Because individuals are constantly trying to minimize energetic costs, we expect behaviors supporting this minimization to be positively valued and trigger automatic reactions. In the current study, we focused on automatic approach tendencies because they are thought to play a key role in the regulation of behaviors (Friese, Hofmann, & Wiers, 2011).

Automatic approach tendencies have been investigated using reaction-time tasks where individuals are instructed to approach or avoid a stimulus as quickly as possible (Cousijn, Goudriaan, & Wiers, 2011; Ernst, et al., 2014; Mogg, Field, & Bradley, 2005; Wiers, et al., 2014; Zhou, et al., 2012). Automatic approach tendencies toward stimuli depicting physical activity and sedentary behaviors have been shown to positively and negatively predict physical activity, respectively (Cheval, et al., 2015; Cheval, et al., 2014). In addition, these studies showed a higher tendency to approach rather than avoid stimuli depicting physical activity and vice versa with sedentary behaviors (Cheval, et al., 2015; Cheval, et al., 2014; Cheval, Sarrazin, Pelletier, & Friese, 2016), thereby suggesting that automatic processes support physical activity. These behavioral results seem inconsistent with our hypothesis stating that behaviors supporting cost minimization activate automatic processes counteracting the implementation of physically active behaviors. However, these higher tendencies to approach physical activity and avoid sedentary behaviors fail to explain the exercise paradox. Behavioral outcomes (i.e., differences in reaction times) fail to provide a complete picture of the processes underlying automatic behaviors in exercise. Behavioral performance may not solely result from facilitation processes but from the competition of both facilitation and inhibition processes in the brain. Investigating the brain correlates of these reaction-time differences is necessary to understand this discrepancy between theory and observed behaviors.

Electroencephalography (EEG) provides the millisecond-range resolution required to capture the brain activity underlying the reaction-time differences used to investigate automatic approach and avoidance tendencies. Lateralized Readiness Potentials (LRP) are used to capture the chronometry of the brain processes underlying an action (Gratton, Coles, Sirevaag, Eriksen, & Donchin, 1988; Smulders & Miller, 2012). Stimulus-locked LRP (S–LRP) reflect sensory integration and response-locked LRP (R–LRP) reflect the subsequent processes involved in motor preparation. Event-Related Potentials (ERP) can reveal brain resources involved in a behavior. Particularly, P1 reflects the automatic allocation of attention toward relevant emotional stimuli (Keus, Jenks, & Schwarz, 2005; Olofsson, Nordin, Sequeira, & Polich, 2008; Smith, Cacioppo, Larsen, & Chartrand, 2003), early N1 reflects conflict monitoring (Botvinick, Cohen, & Carter, 2004; Kerns, et al., 2004; van Veen, Cohen, Botvinick, Stenger, & Carter, 2001), late N1 reflects enhanced perceptual processing during conflict (Ernst, et al., 2013; Kirmizi-Alsan, et al., 2006; Vogel & Luck, 2000), and N2 reflects the inhibition of automatic reactions (Folstein & Van Petten, 2008; van Boxtel, van der Molen, Jennings, & Brunia, 2001).

Here, we investigated the brain regions associated with automatic approach and avoidance reactions toward physical activity and sedentary behaviors. We hypothesized a stronger tendency to approach physical activity than sedentary behaviors and to avoid sedentary behaviors than physical activity (Hypothesis 1a). We expected these tendencies to be stronger in individuals who successfully implement their intention to be physically active (Hypothesis 1b). We further hypothesized that this effect of stimuli on reaction time results from altered processes during sensory integration of these visual stimuli, not during motor preparation. Specifically, we hypothesized that in individual intending to be physically active, like all participants of the current study, sensory integration is shorter (i.e., larger positive deflection and earlier LRP onset latency) when they are asked to approach physical activity and avoid sedentary behaviors compared to approach sedentary behaviors and avoid physical activity (Hypothesis 2). Additionally, consistent with recent conceptual and review articles suggesting that individuals tend to save energy and avoid unnecessary physical exertion (Cheval, et al., 2018; Lee, Emerson, & Williams, 2016; Lieberman, 2015), lower reaction times when approaching physical activity and avoiding sedentary behaviors should require more cortical resources. Accordingly, we hypothesized higher attentional processing (larger P1 and late N1 amplitudes), conflict monitoring (larger early N1 and late N1 amplitudes), and inhibition (larger N2 amplitude) when approaching physical activity compared to sedentary behaviors and when avoiding sedentary behaviors compared to physical activity (Hypothesis 3). We expected these cortical outcomes to be more pronounced in individuals who successfully implement their intention to be physically active (Hypothesis 4).

## 2. Materials and Methods

### 2.1. Participants

Participants were invited to take part in the study through posters in the university. To be included in the study, participants had to be right-handed according to the Edinburgh Handedness Inventory (Oldfield, 1971) and in the preparation (i.e., low level of physical activity with a strong intention to start) or maintenance stage of physical activity (i.e., high level of physical activity for at least 6 months) according to the stage of change questionnaire for exercise behavior (Marcus, Selby, Niaura, & Rossi, 1992). This state-of-change measure was used to ensure that participants were physically active or involved in the process of changing their exercise behavior toward a physically active one. Participants with a history of psychiatric, neurological, or severe mental disorders, or taking psychotropic medication or illicit drugs at the time of the study were excluded. Thirty-seven young volunteers met the eligibility criteria. Eight participants were removed from the analyses due to e-prime and EEG data recording malfunctions resulting in a final sample of 29 participants (age = 22.8 ± 3.0 years; 16 females; body mass index = 21.8 ± 3.1 kg/m^2^) including 14 physically active participants (i.e., maintenance stage) and 15 physically inactive participants with the intention of becoming physically active (i.e., preparation stage). All participants received a 20 CHF voucher. The University of Geneva Ethics committee approved this research and informed consent process.

### 2.2. Pilot Studies

#### 2.2.1. Pilot Study 1: Contextual Stimuli

In Pilot Study 1, we identified the stimuli depicting physical activity and sedentary behaviors to be included in the approach-avoidance task. Thirty-two participants were asked to rate the extent to which 24 stimuli expressed “movement and an active lifestyle” (1 = not at all, 7 = a lot) and “rest and sedentary lifestyle”. To minimize the bias associated with pictures depicting real people, a designer drew pictograms representing physical activity and sedentary behaviors. The size of the stimuli was 200 × 250 pixels. For each stimulus, the “rest and sedentary lifestyle” score was subtracted from the “movement and active lifestyle” score. The 5 stimuli with the largest positive and negative differences were chosen as the stimuli depicting physical activity and sedentary behaviors in the main experiment, respectively. Statistical analyses confirmed that the 5 stimuli depicting physical activity showed higher physical activity scores (M = 5.97, SD = 0.88) than sedentary scores (M = 1.85, SD = 0.69, *t*_(31)_ = −15.33, *p* < 0.001) and that the 5 stimuli depicting sedentary behaviors showed higher sedentary scores (M = 5.30, SD = 1.02) than the physical activity scores (M = 2.15, SD = 0.89, *t*_(31)_ = −10.23, *p* < 0.001).

#### 2.2.2. Pilot Study 2: Neutral Stimuli

In Pilot Study 2, we tested the effect of the neutral stimuli. Thirty-nine participants were asked to rate the extent to which 30 stimuli expressed “rest and sedentary lifestyle” versus “movement and active lifestyle” on a 7-point bipolar response scale (i.e., −3 to +3). We used the 10 stimuli selected in Pilot Study 1 and 20 neutral stimuli based on squares and circles (Figure 1). Statistical analyses confirmed a significant effect of the type of stimulus (i.e., physical activity vs. sedentary behaviors vs. neutral, *F*_(2, 76)_ = 658.14, *p* < 0.001). As expected, post-hoc analyses showed that stimuli depicting physical activity were more strongly related to “movement and active lifestyle” than neutral stimuli (M = 4.66 vs. 2.46, *p* < 0.001), and stimuli depicting sedentary behaviors were more strongly related to “rest and sedentary lifestyle” than neutral stimuli (M = −2.21, *p* < 0.001). Neutral stimuli were not significantly different from a score of zero on the bipolar response scale (*p* = 0.153).

**Figure 1.**
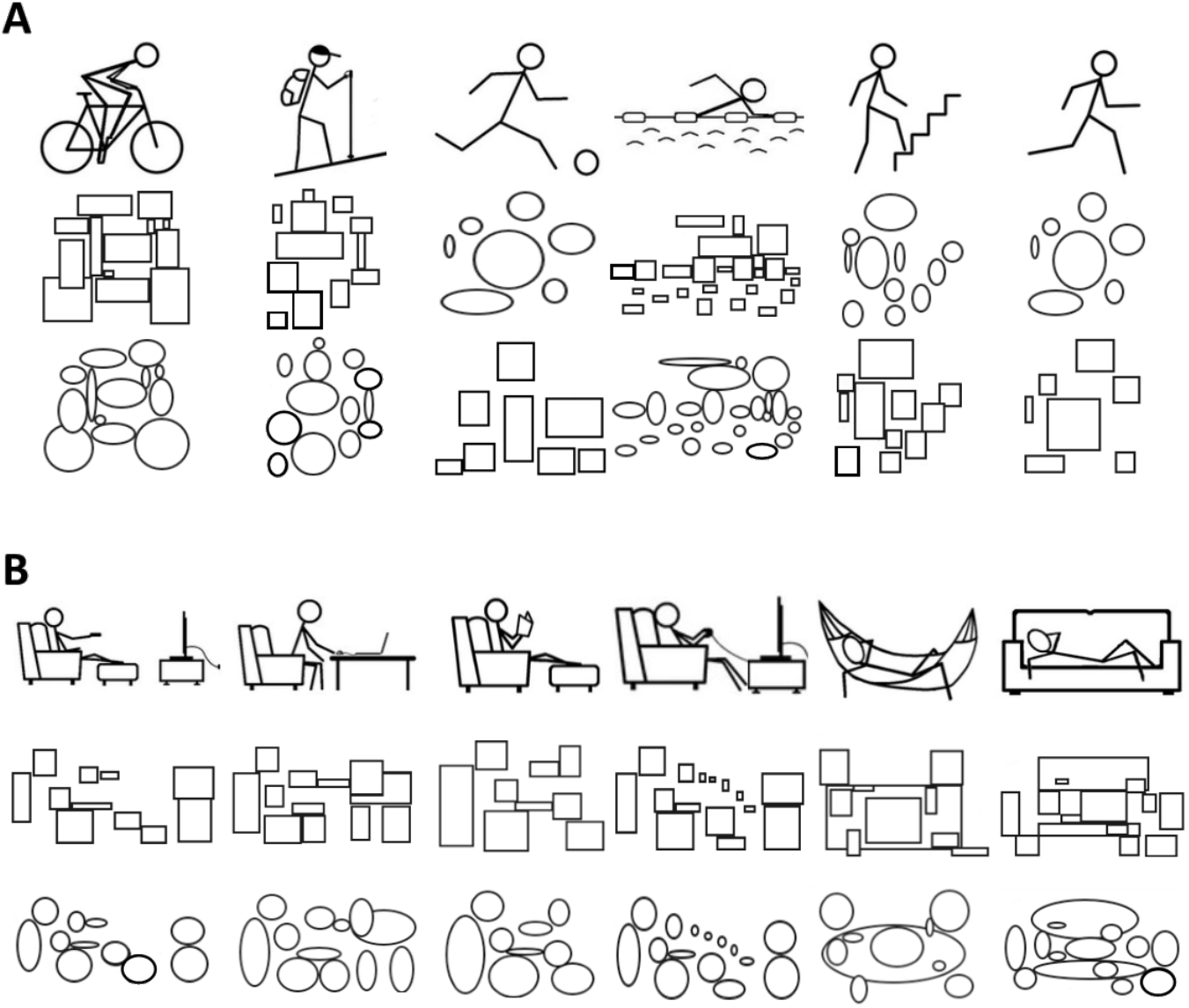
Contextual and neutral stimuli used in the approach-avoidance task. **A.** Stimuli depicting physical activity and neutral stimuli built with circles and squares based on the amount of information (i.e., same number and same size) in the stimuli depicting physical activity. **B.** Images depicting sedentary behaviors and neutral stimuli built with circles and squares based on the amount of information in the stimuli depicting sedentary behaviors. These stimuli were selected based on the results of Pilot Study 1 and Pilot Study 2.

### 2.3. Approach-Avoidance Task

A contextual approach-avoidance task was used to measure automatic approach and avoidance tendencies toward physical activity and sedentary behaviors (Cheval, et al., 2015; Figure 1; Cheval, et al., 2014). Participants were asked to move a manikin on the screen “toward” (approach condition) and “away” (avoidance condition) from images depicting physical activity and sedentary behaviors (Figure 1) by pressing keys on a keyboard. Each trial started with a black fixation cross presented randomly for 250 to 750 ms in the center of the screen with a white background. Then, the manikin appeared in the upper or lower half of the screen. Concurrently, a stimulus depicting “movement and active lifestyle” (i.e., physical activity) or “rest and sedentary lifestyle” (i.e., sedentary behavior) was presented in the center of the screen. Participants quickly moved the human figure “toward” a stimulus (approach) depicting physical activity and “away” from a stimulus (avoidance) depicting sedentary behaviors, or vice versa. After seeing the manikin in its new position for 500 ms, the screen was cleared. In case of an incorrect response, an error feedback (i.e., a cross) appeared at the center of the screen.

A neutral approach-avoidance task was used as a control. In this task, the stimuli depicting physical activity and sedentary behaviors were replaced by stimuli with circles or squares matching the number and size of information in the contextual stimuli (Figure 1). Participants were asked to quickly move the manikin “toward” stimuli with circles and “away” from stimuli with squares, or vice versa. For half of the participants, the neutral stimuli with circles were built based on the stimuli depicting physical activity and the neutral stimuli with squares were built based on the stimuli depicting sedentary behaviors. For the other half of participants, it was the opposite. The neutral approach-avoidance task provided the baseline approach and avoidance tendencies of each individual.

**Figure 2.**
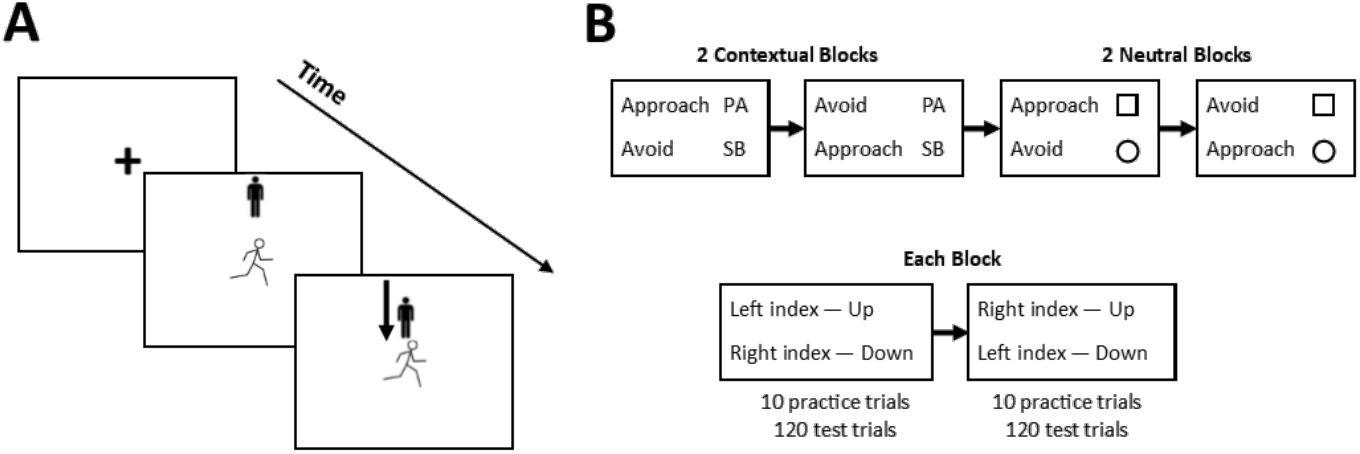
Approach-avoidance task and procedures. **A. Approach-avoidance task.** Trial where the participant is asked to approach a stimulus depicting physical activity. **B. Procedures.** Description of the procedure of the approach-avoidance task. The contextual and the neutral approach-avoidance task, the order of the blocks, and the order of the finger used were counterbalanced across participants. Participants were asked to approach stimuli depicting physical activity (120 trials), avoid stimuli depicting sedentary behaviors (120 trials), avoid stimuli depicting physical activity (120 trials), and approach stimuli depicting sedentary behaviors (120 trials). PA = physical activity; SB = sedentary behaviors.

### 2.4. Experimental Design

Sixty-four participants completed an online questionnaire measuring their stage of change for exercise behavior. This questionnaire was emailed to the participants with a randomly generated identification code. Participants who met the eligibility criteria were invited to the laboratory to sign the informed consent form and respond to the Edinburgh Handedness Inventory (Oldfield, 1971). Then, they sat in front of a computer screen (1280 × 1024 pixels) in a sound-attenuated room, were equipped with EEG recording electrodes, and performed the approach-avoidance task.

The contextual approach-avoidance task was performed in two blocks (Figure 1). In each block, the participants performed 10 practice trials and 240 test trials. During test trials, each of the 10 contextual stimuli appeared 12 times at the top and 12 times at the bottom of the screen. In one block, participants were instructed to approach stimuli depicting physical activity and avoid stimuli depicting sedentary behaviors. In the other block, they were instructed to do the opposite. To compute the LRP, the 240 test trials were divided into two parts. In the first part, participants were asked to press the “8” key with their left index to move the manikin up and the “2” key with their right index to move the manikin down. In the second part, participants were asked to press the “8” key with their right index and the “2” key with their left index. The neutral approach-avoidance task was performed in two additional blocks. The number of practice and test trials was identical as in the contextual approach-avoidance task. In the contextual and neutral approach-avoidance task, the order of the blocks and finger used were counterbalanced across participants, and the stimuli appeared in a random order within each block (Figure 1).

### 2.5. Usual Level of Physical Activity

The usual level of physical activity was assessed using the adapted version of the International Physical Activity Questionnaire (IPAQ; Booth, 2000; Craig, et al., 2003) assessing physical activity undertaken during leisure time during a week. The specific types of activity were classified into 3 categories: Walking, moderate-intensity activities, and vigorous-intensity activities. The usual level of moderate-to-vigorous physical activity (MVPA) in min per week was used in the main analysis.

### 2.6. EEG Acquisition

The electrical signal of the brain was recorded using a 64-channel Biosemi Active-Two system (Amsterdam, Netherlands) with Ag/AgCl electrodes positioned according to the extended 10–20 system. To capture eye movements and blinks, 4 additional flat electrodes were positioned on the outer canthi of the eyes, and above and under the right eye. A reference electrode was positioned on the earlobe. Each active electrode was associated with an impedance value, which was kept below 20 kΩ for each participant. The EEG was continuously recorded with a sampling rate of 1024 Hz.

### 2.7. EEG Processing

Standard processing of EEG data was performed off-line using the Brain Vision Analyzer software, version 2 (Brain Products, Gilching, Germany). Data was down-sampled to 512 Hz. ERPs were segmented from 200 ms prior to 1000 ms after stimulus onset. Electrodes that were noisy over the entire recording (i.e., 2.5% of the electrodes) were interpolated using a spherical spline (Perrin, Pernier, Bertrand, & Echallier, 1989). A baseline correction was applied using the 200 ms prestimulus period. ERPs and LRPs were obtained by averaging the trials for each condition on the data that was filtered with a low-cutoff at 0.1 Hz and a high-cutoff at 30 Hz. Ocular movements and blink correction was performed using the implemented standard algorithm (Gratton, Coles, & Donchin, 1983). Trials with other artefacts were removed using a semi-automatic procedure (amplitude allowed: −100 to +100 μV) resulting in a total of 11% removed trials.

### 2.8. EEG Metrics

#### 2.8.1. Event-Related Potentials

The P1 ERP peaks around 100–130 ms post-stimulus over the lateral occipital electrodes and reflects activity in the extrastriate cortex (Luck, 2014). P1 reflects automatic attention allocation toward relevant emotional stimuli (Keus, et al., 2005; Olofsson, et al., 2008; Smith, et al., 2003). The N1 ERP can be divided in several subcomponents, with earlier effects appearing on anterior electrodes and later effects appearing on posterior electrodes (Luck, 2005; Ernst et al., 2013). The early N1 peaks around 100–150 ms post-stimulus over the anterior electrodes and has been linked to the activity of the anterior cingulate cortex (Mulert, Gallinat, Dorn, Herrmann, & Winterer, 2003; Mulert, et al., 2001). This activity occurs during incentive conditions, with higher incentives leading to higher anterior N1 amplitudes (Mulert, Menzinger, Leicht, Pogarell, & Hegerl, 2005). Moreover, the activity of the anterior cingulate cortex has been linked to conflict monitoring (Botvinick, et al., 2004; Kerns, et al., 2004; van Veen, et al., 2001). The late N1 peaks around 150–200 ms post-stimulus over the posterior electrodes and reveals activity in the lateral occipital cortex (Luck, 2014). This activity is elicited by discriminative processing in spatial attention tasks (Vogel & Luck, 2000) leading to enhanced perceptual processing of relevant stimuli. In the context of approach-avoidance tasks, the late N1 has been elicited in conflict-related conditions (Ernst, et al., 2013; Kirmizi-Alsan, et al., 2006). The fronto-central N2, which peaks around 200–400 ms post-stimulus (Ernst et al., 2013), is thought to reflect inhibition of automatic reactions (Folstein & Van Petten, 2008; van Boxtel, et al., 2001).

#### 2.8.2. Lateralized Readiness Potentials

LRP are movement-related brain potentials reflecting hand-specific motor preparation (Leppänen, Tenhunen, & Hietanen, 2003; Masaki, Takasawa, & Yamazaki, 2000) and can detect subtle activations that do not necessarily lead to an overt motor response (Dehaene, et al., 1998). LRP can be assessed to capture the chronometry of the brain processes underlying an action and to infer the cognitive demand related to this action (Smulders & Miller, 2012). LRP can be divided into two components. The stimulus-locked LRP (i.e., measured with respect to the stimulus onset; S–LRP) reflects sensory integration and the response-locked LRP (i.e., measured with respect to the manual response; R–LRP) reflects the subsequent processes involved in motor preparation (Luck & Kappenman, 2011; Mordkoff & Gianaros, 2000; Rinkenauer, Osman, Ulrich, Müller-Gethmann, & Mattes, 2004). In a choice reaction-time task involving both upper limbs, positive deflections indicate response preparation of the correct limb, whereas negative deflections indicate a short-lived covert activation of the incorrect limb (Dehaene, et al., 1998). In other words, in incongruent conditions (i.e., when the intended response hampers the selection of the required response), the stimulus induces a covert motor activation that mismatches with the overt response required by the task, leading to a competition between responses.

LRPs were computed in each condition using the double subtraction technique. The signal from the electrodes contralateral to the response was averaged in each participant (C3: Left hemisphere and C4: Right hemisphere). Then, the following formula was applied:

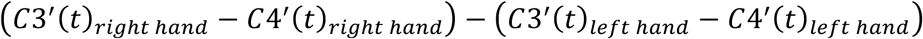

where *C3′*(*t*) and *C4′*(*t*) are the potentials at *C3′* and *C4′* scalp sites, respectively, for multiple time points (Smulders & Miller, 2012). The difference between contralateral and ipsilateral potentials on these electrodes allowed the identification of a specific response (right or left hand) for each condition. For LRPs relative to stimulus onset, epochs were calibrated 200 ms before and 1500 ms after stimulus onset. For LRPs relative to response onset, epochs were calibrated 500 ms before and 100 ms after response onset.

### 2.9. Statistical Analyses

#### 2.9.1. Behavior

Incorrect responses and responses below 150 ms and above 1500 ms were excluded as recommended by Krieglmeyer and Deutsch (2010). The relative reaction times to approach (or avoid) stimuli depicting sedentary behaviors were calculated by subtracting the median reaction time of the participant when approaching (or avoiding) neutral stimuli from each reaction time when approaching (or avoiding) stimuli depicting sedentary behaviors. This subtraction was applied to control for the reaction time associated with the tendency to approach and avoid neutral stimuli. The same procedure was applied to the stimuli depicting physical activity. Behavioral data were analyzed with linear mixed models, which take into account both the nested (multiple measurements within a single individual) and crossed (participants and stimuli) random structure of the data, thereby providing accurate parameter estimates with acceptable type I error rates (Boisgontier & Cheval, 2016). Moreover, linear mixed models avoid data averaging which keeps the variability of the responses in each condition and increases power compared with traditional approaches such as the analysis of variance (ANOVA) (Judd, Westfall, & Kenny, 2017). We built a model using the lme4 and lmerTest packages in the R software (Bates, Mächler, Bolker, & Walker, 2014; Kuznetsova, Brockhoff, & Christensen, 2015) and specified both participants and stimuli as random factors. Action (−0.5 for approach trials; 0.5 for avoidance trials), Stimuli (−0.5 for stimuli depicting physical activity; 0.5 for stimuli depicting sedentary behaviors), the usual level of MVPA (continuous; standardized), and their interactions were included as fixed factors in the model. A random error component was included for Action and Stimuli.

An estimate of the effect size was reported using the conditional pseudo R^2^ computed using the MuMin package of the R software (Barton, 2009). Simples slopes, region of significance, and confidence bands were estimated using Preacher and colleagues’ computational tools for probing interactions in mixed models (Preacher, Curran, & Bauer, 2006). Statistical assumptions associated with linear mixed models, including normality of the residuals, linearity, multicollinearity (variance inflation factors), and undue influence (Cook’s distances) were met.

#### 2.9.2. Event-Related Potentials

Because this study was the first to use ERPs to investigate approach and avoidance reactions to physical activity and sedentary behaviors stimuli, it was not possible to formulate specific a priori hypotheses on the spatiotemporal distribution of the potential effects. Therefore, we performed a whole-scalp analysis (64 electrodes) from 0 (stimulus appearance) to 800 ms using a cluster-mass permutation test (Maris & Oostenveld, 2007), which is appropriate for exploratory analyses and delimiting effect boundaries when little guidance is provided by previous research (Groppe, Urbach, & Kutas, 2011; Luque, et al., 2017; Manly, 1997). To fit the analysis with the experimental design and use resampling methods, we perform F-tests of repeated measures ANOVA and the null distribution was computed using permutations of the reduced residuals (Kherad-Pajouh & Renaud, 2015). The family-wise error rate was controlled using the cluster-mass test (Maris & Oostenveld, 2007) with a threshold set at the 95th percentile of the F statistic. For the cluster-mass test, we defined the spatial neighborhoods between electrodes using an adjacency matrix. Each pair of electrodes with a Euclidian distance smaller than delta = 35mm was defined as adjacent, where delta is the smallest value such that the graph created by the adjacency matrix is connected.

#### 2.9.3. Lateralized Readiness Potentials

The amplitude LRP were analyzed with a 2 (Action: approach vs. avoidance) × 2 (Stimuli: physical activity vs. sedentary behaviors) × the usual level of MVPA (continuous) mixed-subject design analysis of variance (ANOVA). LRP outcomes were analyzed using ANOVA because the use of linear mixed models has not been implemented for LRP analyses yet. We used the relative signal, i.e., the difference between the amplitude of the contextual and neutral stimuli. The LRP onsets were measured and analyzed by applying the jackknife-based procedure (Ulrich & Miller, 2001). LRP onset measures were submitted to ANOVA with F-values corrected as follows:

*Fc* = *F*/(*n* – 1)^2^, where *Fc* is the corrected F-value and *n* the number of participants (Ulrich & Miller, 2001). The Greenhouse-Geisser epsilon correction was applied to adjust the degrees of freedom of the F-ratio when appropriate. LRP measurements (amplitude and onset latencies) were computed based on the average of left and right manual responses, with respect to the experimental condition.

#### 2.9.4. Sensitivity Analyses

To examine the robustness of the simple effects of approaching versus avoiding sedentary behaviors and physical activity stimuli, we performed three sensitivity analyses: using only circle-based neutral stimuli, using only square-based neutral stimuli, and replacing the usual level of physical activity by the stage of change for exercise.

### 2.10. Data and Code Accessibility

All data and code are available in Zenodo (doi: 10.5281/zenodo.1169140).

## 3. Results

### 3.1. Descriptive Results

Results showed that participants in the preparation stage self-reported lower usual level of physical activity than participants in the maintenance stage of physical activity (93.6 ± 74.0 vs. 330.0 ± 160.0 minutes per week, *p* < 0.001). Body mass index (22.3 ± 3.7 vs. 21.3 ± 2.3 kg/m^2^, *p* = 0.415), age (23.4 ± 3.1 vs. 22.2 ± 2.9 years, *p* = 0.276), and sex (8 females and 6 males vs. 7 males and 8 females, *p* = 0.999) were not significantly different across groups. Figure 3 reports reaction times to approach and avoid stimuli depicting physical activity, sedentary behaviors, and neutral stimuli in more (Figure 3A) and less physically active participants (Figure 3B).

**Figure 3.**
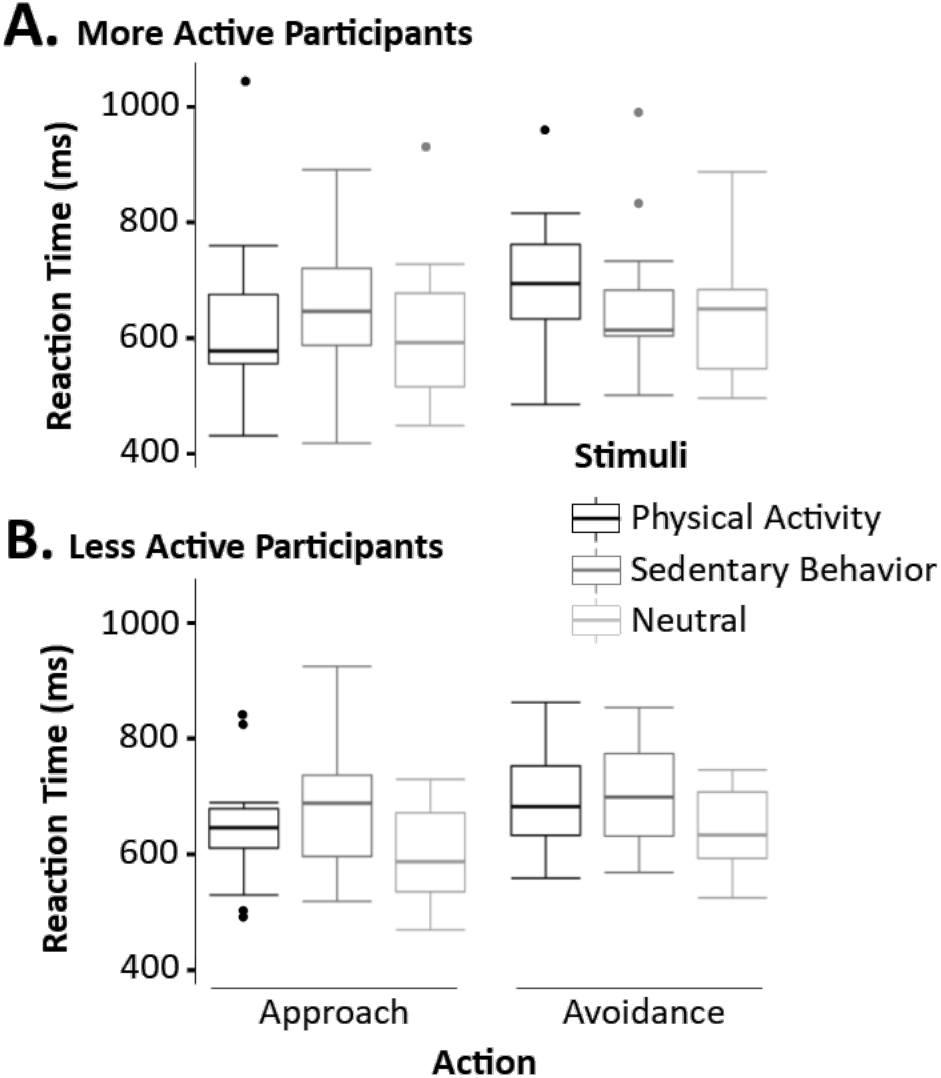
Descriptive results showing reaction time to approach and avoid stimuli depicting physical activity, sedentary, and neutral stimuli in less (A) and more physically active participants (B). Groups were determined by a mid-point split of the moderate-to-vigorous physical activity variable. The middle of the boxplot = median, lower hinge = 25% quantile, upper hinge = 75% quantile, lower whisker = smallest observation greater than or equal to lower hinge – 1.5 × interquartile range, upper whisker = largest observation less than or equal to upper hinge + 1.5 × interquartile range.

### 3.2. Behavioral Results

Results of the linear mixed models (Table 1) showed no significant main effects of action (*p* = 0.973), stimuli (*p* = 0.426), and usual level of MVPA (*p* = 0.295). However, the two-way interaction between action and stimuli was significant (b = −63.23, *p* < 0.001). Simple effect tests showed that participants approached stimuli depicting physical activity faster than sedentary behaviors (b = 37.74, *p* < 0.001). Conversely, participants were slower at avoiding physical activity than sedentary stimuli (b = −25.50, *p* = 0.006). Additionally, results showed that participants were faster at approaching than avoiding physical activity (b = 31.42, *p* < 0.001), whereas they were faster at avoiding than approaching sedentary behaviors (b = −31.82, *p* < 0.001). The three-way interaction between action, stimuli, and MVPA for exercise was significant (b = −38.50, *p* < 0.001). As illustrated in Figure 4, results showed that the two-way interaction between action and stimuli was significantly more pronounced when the usual level of MPVA was high (+1SD; b = −101.81, *p* < 0.001) than low (−1SD; b = −24.80, *p* = 0.006). In this model, the variables under consideration explained 14.9% of the variance in reaction time. The region of significance of the simple slope showed that participants were slower at approaching sedentary than physical activity stimuli. This effect was more pronounced when MVPA was higher (Figure 4A, upper panel). For example, when MPVA was low (−1SD), participants took ~20 ms longer to approach sedentary than physical activity stimuli (b = 20.456, *p* = 0.0382; Figure 4B, lower panel). When MPVA was high (+1SD), participants took ~55 ms longer (b = 55.19, *p* < 0.001; Figure 4B, upper panel). The region of significance started at a lower MPVA in the approach than avoidance condition of sedentary behaviors (lower bound at MVPA = −0.47).

**Table 1.**
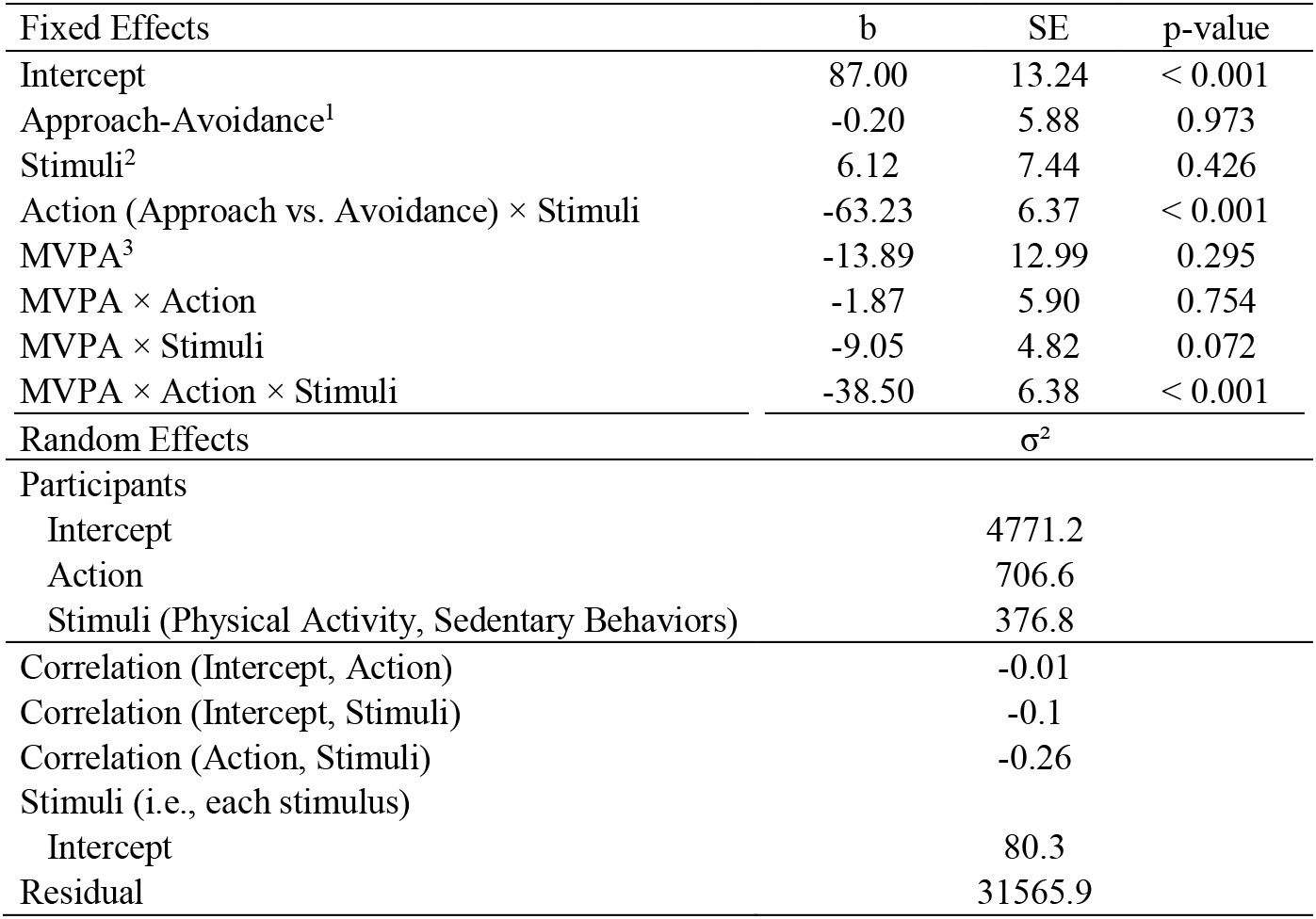
Results of the linear mixed models predicting the relative reaction time required to approach and avoid stimuli depicting physical activity and sedentary behaviors as a function of the usual level of moderate-to-vigorous physical activity (MVPA). The relative reaction time to approach (avoid) stimuli associated with physical activity and sedentary behaviors compared to neutral stimuli was obtained by subtracting each participant’s median reaction times to approach (avoid) neutral stimuli from each specific reaction time to approach (avoid) stimuli depicting physical activity and sedentary behaviors; ^1^ −0.5 = approach; 0.5 = avoidance; ^2^ −0.5 = physical activity; 0.5 = sedentary behaviors; ^3^ continuous; SE = standard error.

**Figure 4.**
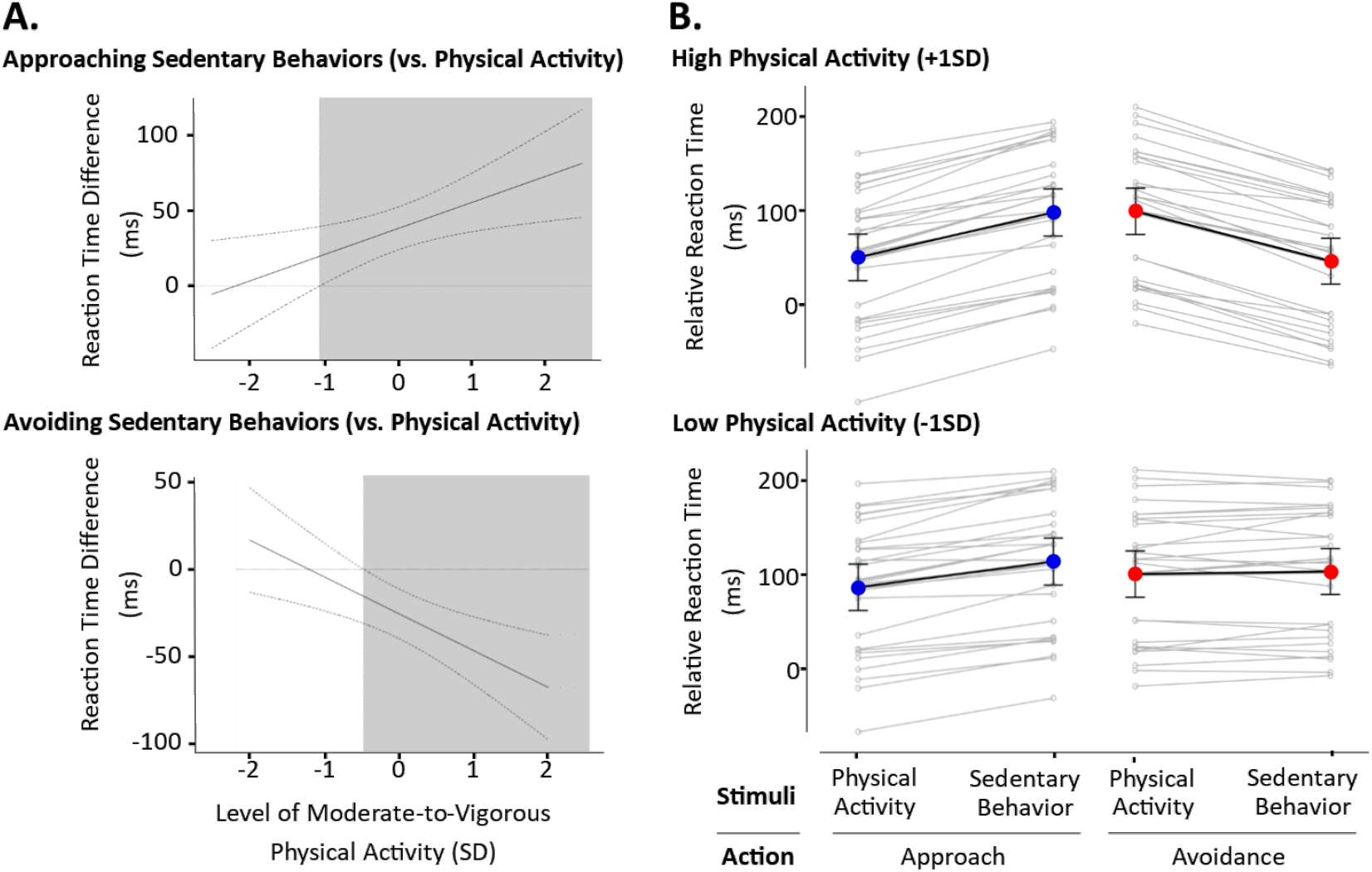
Results of the linear mixed models. **A. Region of significance of the effect to approach (upper left panel) and avoid (lower left panel) sedentary behaviors relative to physical activity as a function of MVPA.** A positive difference indicates a slower reaction time to approach (upper left panel) and avoid (lower left panel) sedentary behaviors relative to physical activity. SD = standard deviation; solid line = mean; dashed line = 95% confidence interval; grey area = region of significance *(p* < 0.05). **B. Relative mean reaction time in ms as predicted by the linear mixed model to approach (blue dot) and avoid (red dot) physical activity and sedentary behaviors at low (−1SD) (upper right panel) and (+1SD) high level of MVPA (lower right panel).** Grey dots represent each individual’s mean of the repeated trials for each condition (Action and Stimuli). Errors bars represent range going from −1.96 SD to +1.96 SD for each condition.

The region of significance of the simple slope also showed that participants were faster at avoiding sedentary than physical activity stimuli. This effect was more pronounced when MVPA was high (Figure 4A, lower panel). For example, when MPVA was low (−1SD), reaction times were similar when participants avoided sedentary and physical activity stimuli (b = −4.35, *p* = 0.661, Figure 4B, lower panel). However, when MPVA was high (+1SD), participants were ~46 ms faster when avoiding sedentary compared to physical activity stimuli (b = −46.62, *p* < 0.001; Figure 4B, upper panel).

### 3.3. Lateralized Readiness Potentials

#### 3.3.1. S–LRP Onset Latency

Results of the S–LRP onset latency did not show a significant main effect of action (*p_c_* = 0.96) and usual level of MVPA (*p_c_* = 0.96). However, results showed a significant main effect of stimuli (*F*_(1, 27)_ = 5310.33, *p* < 0.001, partial *η*^2^ = 0.99, *F*_c(1, 27)_ = 6.73) and a significant interaction between action and stimuli (*F*_(1, 27)_ = 9015.19, *p* < 0.001, partial *η*^2^ = 0.97, *F*_c(1, 27)_ = 12.44; Figure 5). Simple test effects revealed a longer onset latency to approach sedentary (32 ms) than physical activity stimuli (−18 ms, *p_c_s* < 0.001). Conversely, no significant differences emerged between avoiding sedentary and physical activity stimuli, and between approaching and avoiding sedentary (*p_c_* = 0.320) or physical activity stimuli (*p_c_* = 0.640). All the other effects were nonsignificant.

**Figure 5.**
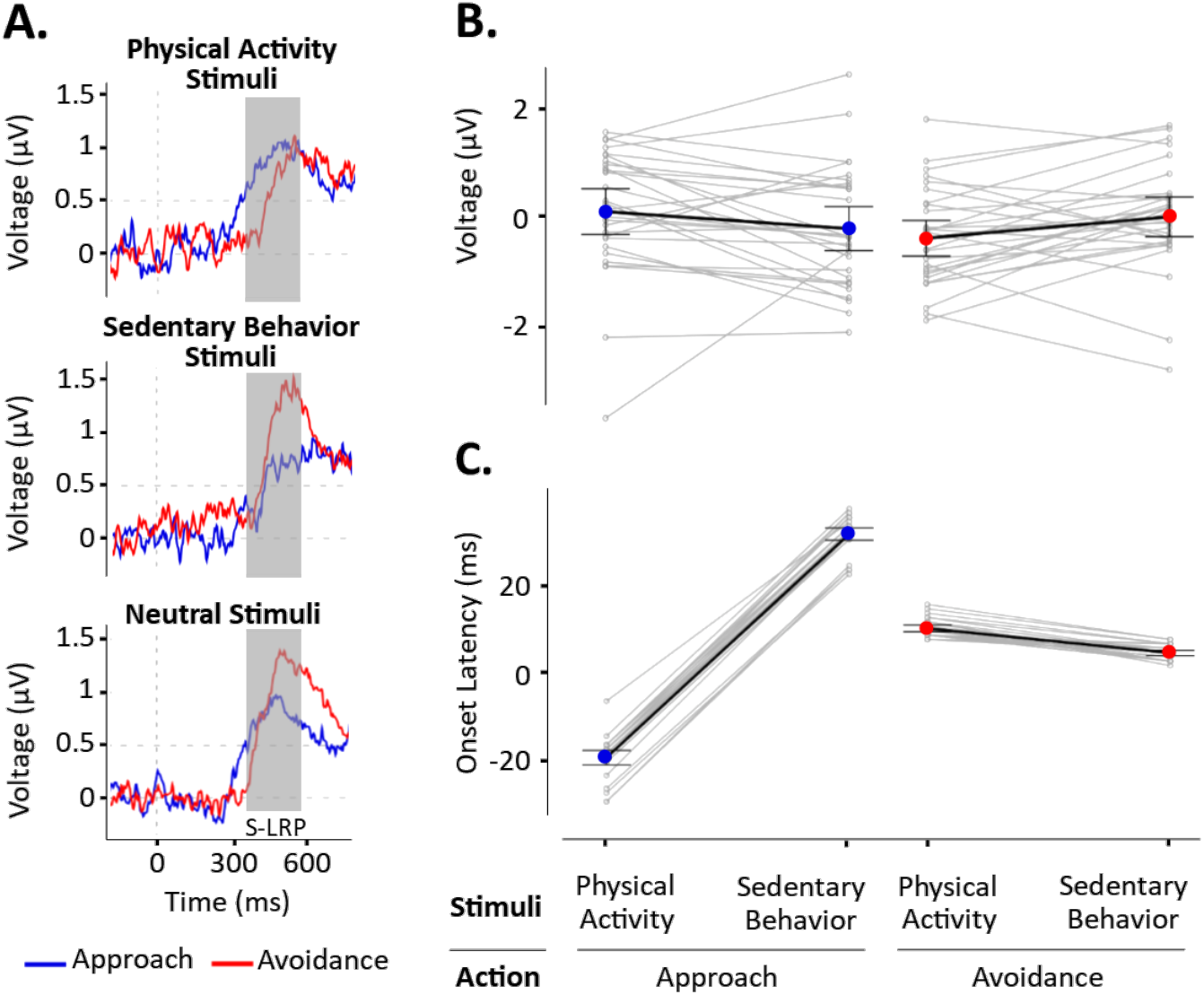
S–LRP results. **A.** Lateralized Readiness Potential (LRP) signal in the 200–800 ms range when approaching (blue line) and avoiding (red line) stimuli depicting physical activity, sedentary behaviors, and neutral stimuli. The grey area represents the range of time associated with the stimulus-locked LRP (S–LRP). **B.** S–LRP amplitudes when approaching (blue dot) and avoiding (red dot) stimuli depicting physical activity and sedentary behaviors. The amplitudes reported here represents amplitudes associated with contextual stimuli (i.e., depicting physical activity or sedentary behaviors) relative to the amplitudes associated with neutral stimuli. Accordingly, a positive amplitude represents a larger positive deflection associated with the contextual stimuli compared to the neutral stimuli. **C.** S–LRP onset latencies when approaching (blue dot) and avoiding (red dot) stimuli depicting physical activity and sedentary behaviors. The onset latencies reported here were relative to the onset latencies associated with neutral stimuli. A negative onset latency represents a shorter onset latency in the contextual than neutral stimuli. It should be noted the jackknife procedure requires to apply the Greenhouse-Geisser epsilon correction to adjust the degrees of freedom of the F-ratio. It should also be noted that the S–LRP amplitudes showed three individuals that may appear as extremes. However, the potential extreme values were going in the opposite direction as the observed effect. Therefore, the effect was significant despite these individuals, and not because of them.

#### 3.3.2. S–LRP amplitude

The mean S–LRP amplitude was measured within the 385–580 ms range, where the overall S–LRP was maximal. Results of the mixed-subject design ANOVA showed non-significant main effects of action (*p* = 0.469), stimuli (*p* = 0.622), and usual level of MVPA (*p* = 160). However, results showed a significant interaction between action and stimuli (*F*_(1, 27)_ = 4.86, *p* = 0.036, partial η^2^ = 0.152; Figure 5). Simple test effects revealed that the avoidance of stimuli depicting physical activity (−0.35 μV, SE = 0.16) elicited a larger negative deflection than the avoidance of stimuli depicting sedentary behaviors (0.041 μV, SE = 0.18, *t*_(28)_ = −2.34, *p* < 0.026) and the approach of physical activity (0.14 μV, SE = 0.21, *t*_(28)_ = −2.10, *p* < 0.04). The other simple effects were not significant (*ps* > 0.127). All the other effects were nonsignificant.

#### 3.3.3. R–LRP

The grand average waveforms of R–LRP are shown in Figure 5. The mean amplitude of R–LRP was measured within the –352 to –60 ms range, where its overall amplitude was maximal for the following negative deflection. Results of the mixed-subject design ANOVA did not show significant main effects of action (*p* = 0.873), stimuli (*p* = 0.220), and usual level of MVPA (*p* = 0.180). The two and three-way interactions were also not significant (*p_s_* > 0.263). In line with the results of the R–LRP amplitudes, results of the mixed-subject design ANOVA testing the R–LRP onset latency showed nonsignificant main effects of action (*p_c_* = 0.882), stimuli (*p_c_* = 0.718), and usual level of MVPA (p_c_ = 0.546). The two and three-way interactions were also not significant (*p_c_s* > 0.985).

### 3.4. Event-Related Potentials

#### 3.4.1. Cluster-Mass Analysis

Results of the cluster-mass analysis showed a significant main effect of stimuli at several time-points in the 100–630 ms range (*p* = 0.0002) with a more negative amplitude for stimuli depicting sedentary behaviors compared to stimuli depicting physical activity. This effect was particularly pronounced and spread between 150 and 350 ms (Figure 6A). The main effect of action was not significant. Results also showed a significant two-way interaction between action and stimuli at several time points between 100 and 400 ms in an area including frontal, central, and parietal sites (*p* = 0.0190; Figure 6C). This interaction effect was particularly pronounced and spread in the 150–325 ms range. Figure 7 illustrates the topographical map for this range period for each condition. Simple effect tests revealed significant amplitude differences when avoiding sedentary behaviors versus physical activity (*p* = 0.0002), when approaching sedentary behaviors versus physical activity (two clusters showed significant effects with *p* = 0.0168 and p = 0.0002), and when avoiding versus approaching sedentary behaviors (*p* = 0.0246). Results showed no significant differences when avoiding or approaching physical activity (lowest *p* = 0.0956). The three-way interaction between action, stimuli, and MVPA was not significant.

**Figure 6.**
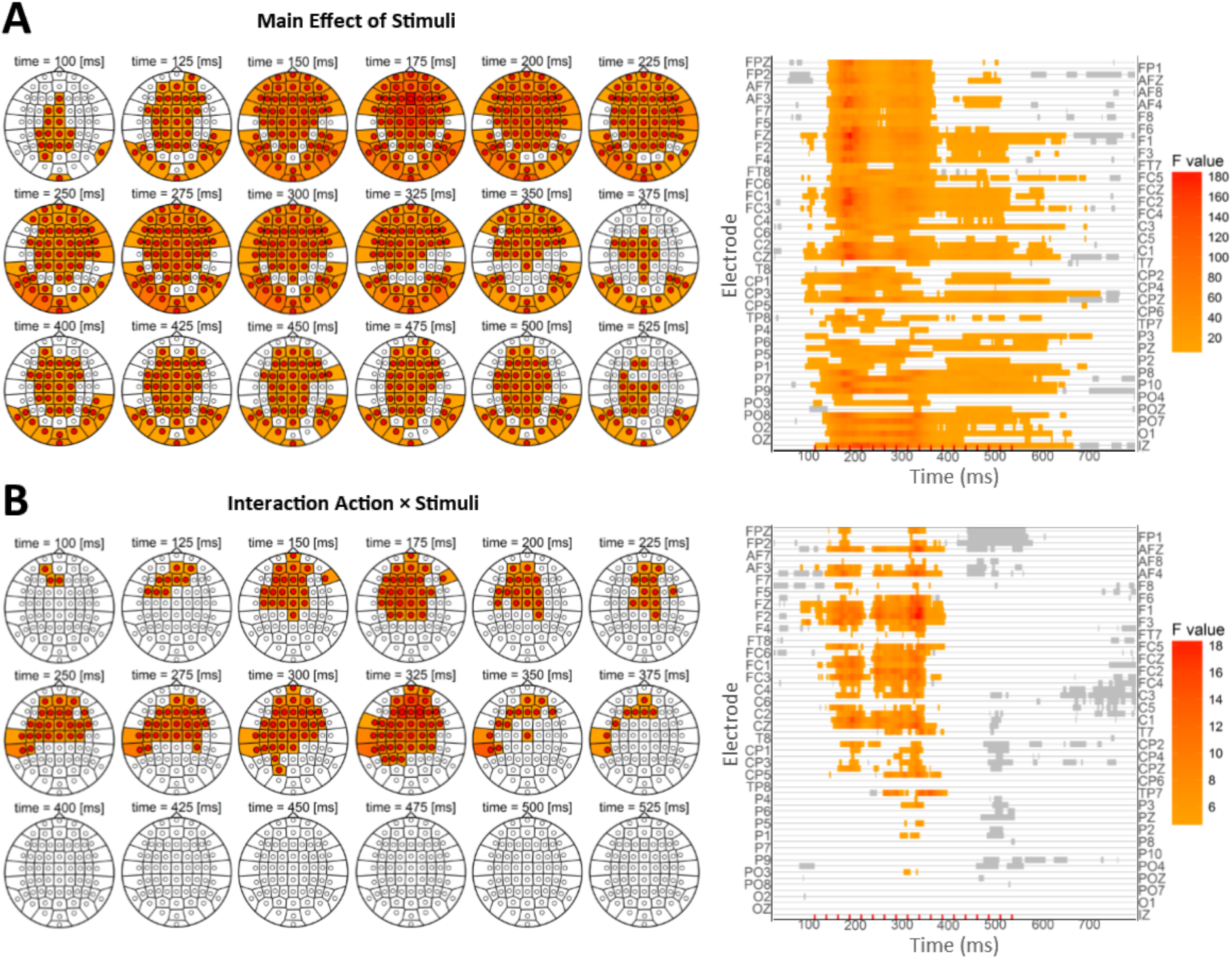
ERP results of the whole-scalp analysis. **A.** Main effect of stimuli for all the electrodes in the 0–800 ms range. **B.** Two-way interaction between action and stimuli for all electrodes in the 0–800 ms range. Results were based on a cluster-mass analysis using non-parametric permutation test and using the family-wise error rate correction.

#### 3.4.2. P1, N1, and N2 ERPs

The first effect, within the 80–130 ms range, was compatible with the P1 ERP and was qualified by a main effect of stimuli with a more positive amplitude for sedentary than physical activity stimuli (Figure 8 illustrates results in P9).

The second effect, within the 100–150 ms range, was compatible with the early N1 ERP and was qualified by a main effect of stimuli with a more negative amplitude for stimuli depicting sedentary behaviors compared to physical activity. Moreover, a two-way interaction between action and stimuli emerged at the end of the period. This interaction was characterized by a more negative amplitude for avoiding sedentary behaviors compared to physical activity. This simple effect also emerged in the approach condition but was less pronounced and emerged at the end of the time period only. Results revealed a more negative amplitude for avoiding compared to approaching sedentary behaviors and a more negative amplitude for approaching compared to avoiding physical activity. However, these simple effects were not significant (Figure 7 illustrates results from this analysis with the Fcz electrode).

**Figure 7.**
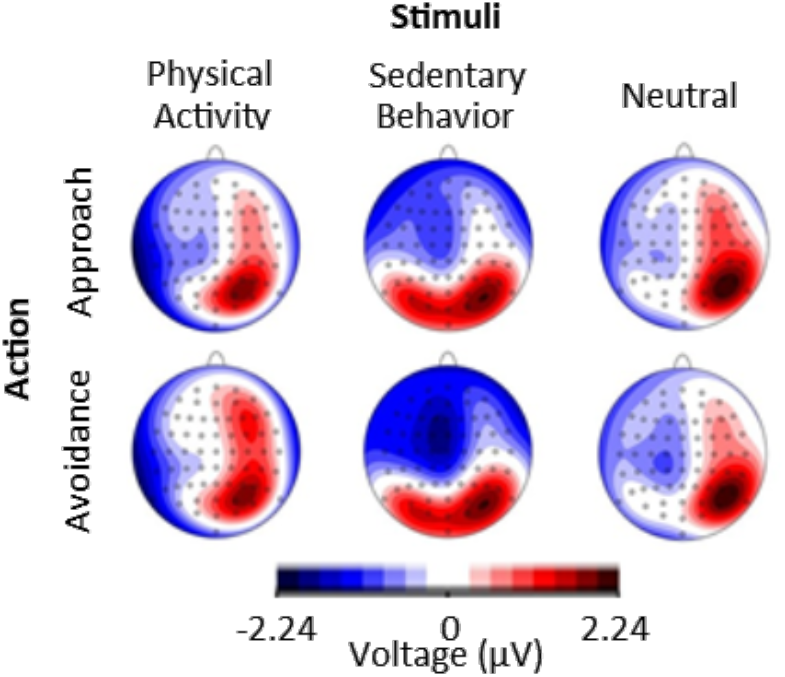
Topographical figure mapping the differences between each condition in the 150–325 ms. The 150–325 ms range was chosen because the interaction between action and stimuli was particularly pronounced and spread within this range.

The third effect, within the 150–180 ms range, was compatible with the late N1 ERP and was qualified by a main effect of stimuli with a more negative EEG amplitude for stimuli depicting sedentary behaviors compared to physical activity (Figure 8 illustrates results in P9).

**Figure 8.**
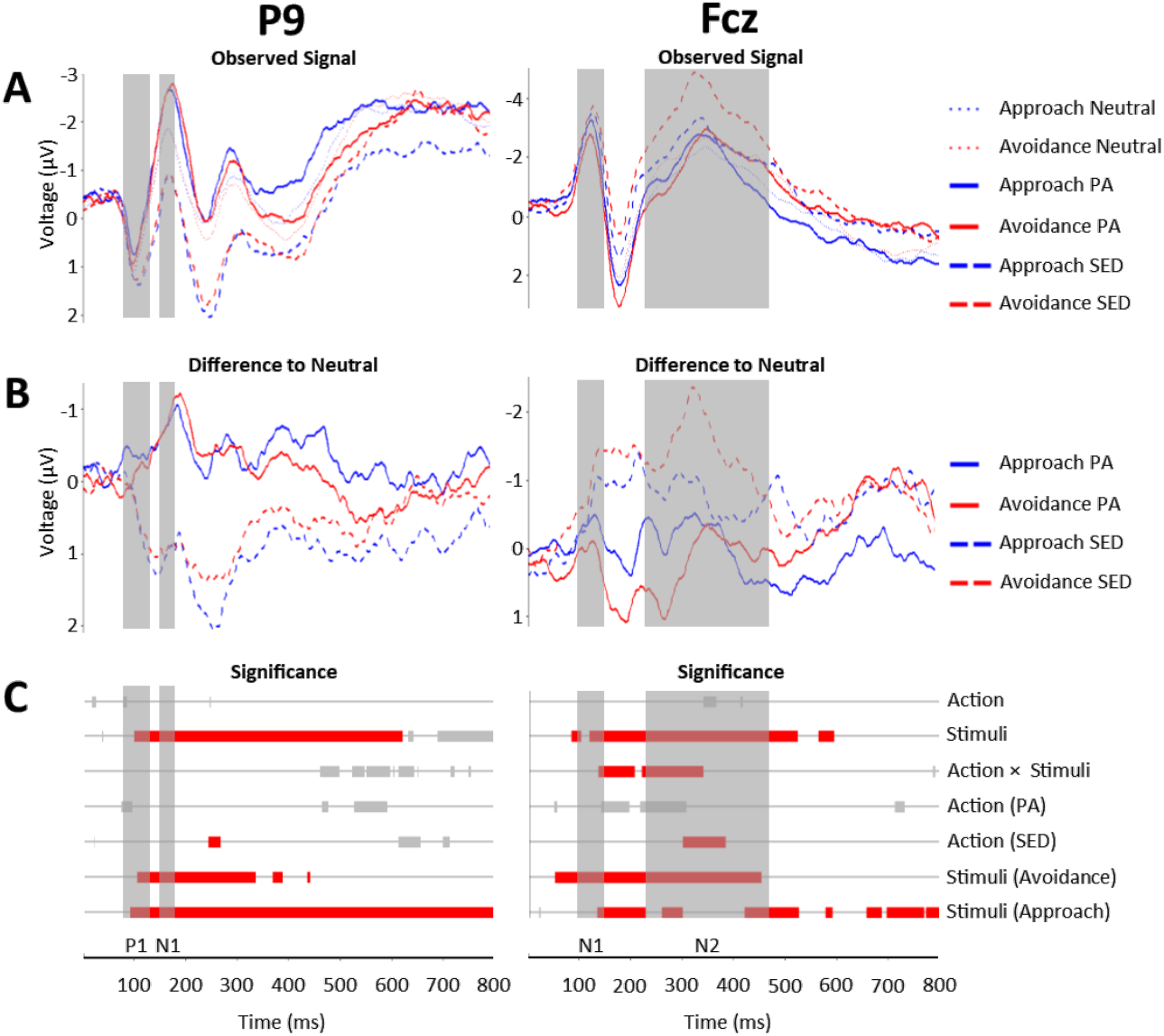
ERP results. **A.** Observed ERP signal in the 0–800 ms for all the conditions. **B.** Difference in the observed ERP signal for approaching and avoiding stimuli depicting physical activity and sedentary behaviors relative to the observed ERP signal for approaching and avoiding neutral stimuli. **C.** Significant effects after the familywise error rate correction. Red bars represent significant effects. Grey bars represent significant effects that did not survive the familywise error correction. For the electrode P9, the first grey area (80–130 ms range) corresponds to P1 and the second grey area (150–180 ms range) represents late N1. For the electrode Fcz, the first grey area (100–150 ms range) represents early N1 results and the second grey area (230–470 ms range) represents N2.

The fourth effect, within the 230–470 ms range, was compatible with the N2 ERP and was qualified by a main effect of stimuli, with a more negative amplitude for stimuli depicting sedentary behaviors compared to physical activity. Moreover, the N2 ERP was qualified by a two-way interaction between action and stimuli. This interaction was characterized by a more negative amplitude for avoiding sedentary behaviors compared to physical activity. This simple effect also emerged for physical activity but was less pronounced and not significant during the whole period. Additionally, results revealed a more negative amplitude for avoiding compared to approaching sedentary behaviors but a more negative amplitude for approaching compared to avoiding physical activity. However, only the simple effect of action for sedentary behaviors was significant (Figure 8 illustrates results in Fcz). These P9 and Fcz ERP outcomes were illustrated as they best represented the observed effects in terms of effect sizes. Moreover, they are traditionally used to index the respective ERPs in the literature.

### 3.5. Sensitivity Results

Overall, the behavioral results of the sensitivity analyses were consistent with the main results, except for the simple effects of approaching versus avoiding stimuli depicting physical activity and sedentary behaviors, which were dependent on the type of neutral stimuli (i.e., based on circles vs. squares). Overall, the LRP results of the sensitivity analyses were consistent with the main results. As for the main analysis, the stage of change for exercise did not modulate the effects on R–LRP amplitudes, S–LRP amplitudes, and onsets. Overall, the ERP results of the sensitivity analyses were consistent with the main results, except for the simple effect of approaching versus avoiding stimuli depicting sedentary behaviors, which did not survive the error rate correction when using either circles or squares as neutral stimuli.

## 4. Discussion

This study revealed that the brain processes underlying faster reactions to approach physical activity and avoid sedentary behaviors occur during sensory integration (larger positive deflection and earlier S–LRP onset latency), not during motor preparation (no effect on the R– LRP components). Results also showed, for the first time, that avoiding sedentary behaviors triggers higher conflict monitoring (larger early N1), and inhibition (larger N2) than avoiding physical activity, irrespective of the usual level of MVPA. These findings suggested that higher levels of control were required to counteract a general trend to approach sedentary behaviors. In line with the affective-reflective theory recently proposed by Brand and Ekkekakis (2018), these results suggest that exercise behavior could be the product of interactions between driving and restraining forces toward physical activity and sedentary behaviors. These interactions challenge the mainstream multidimensional theorizing of exercise behavior with physical activity and sedentary behaviors being conceived as two independent behaviors with different psychological roots. Our results support a unidimensional conception of exercise behavior positioning sedentary behaviors and physical activity on the same continuum of behaviors involving similar psychological processes.

### 4.1. Behavioral Outcomes

#### 4.1.1. Approach and Avoidance Tendencies

Results showed that participants were faster at approaching stimuli depicting physical activity compared to sedentary behaviors, whereas they were faster at avoiding stimuli depicting sedentary behaviors compared to physical activity (Hypothesis 1a). Moreover, results showed that these behavioral outcomes were more pronounced when the usual level of MVPA was higher (Hypothesis 1b). Particularly, results suggested that avoiding sedentary behaviors was more difficult in less physically active individuals. These findings are consistent with previous studies suggesting that automatic reactions toward sedentary behaviors play an important role in the regulation of physical activity (Brand & Ekkekakis, 2018; Cheval, et al., 2015; Cheval, et al., 2014).

#### 4.1.2. Approach Bias

Additionally, previous behavioral studies showed that young and middle-aged adults, especially those who are physically active, exhibited a positive approach bias toward stimuli depicting physical activity (i.e., they were faster at approaching compared to avoiding physical activity stimuli), but a negative approach bias toward sedentary behaviors (i.e., they were faster at avoiding compared to approaching sedentary behaviors) (Cheval, et al., 2015; Cheval, et al., 2014; Cheval, Sarrazin, Pelletier, et al., 2016). However, these previous experiments did not control for the tendency to approach or avoid neutral stimuli. Yet, some individuals may have a tendency to approach rather than avoid neutral stimuli (i.e., a general approach bias), whereas others may have a tendency to avoid rather than approach neutral stimuli (i.e., a general avoidance bias). As such, this absence of control for neutral stimuli may have biased the results. For the first time, our study examined the approach and avoidance tendencies toward stimuli depicting physical activity and sedentary behaviors relative to neutral stimuli. Results showed faster approach than avoidance of physical activity and the opposite for sedentary behaviors. These effects were more pronounced when the usual level of MVPA was higher. These findings are consistent with the suggestion that physically active individuals may have developed positive affective association with physical activity (Brand & Ekkekakis, 2018; Williams, et al., 2008) and/or efficient strategies to increase their automatic tendencies to approach physical activity and decrease those to avoid sedentary behaviors.

### 4.2. Cortical Outcomes

The behavioral results reported in the previous section are inconsistent with the fact that most individuals fail to exercise regularly despite the intention to be physically active (Rhodes & Bruijn, 2013; Rhodes & Dickau, 2012). Therefore, investigating the brain correlates of these reaction-time differences was necessary to understand this discrepancy. The current study examined for the first time the cortical activity associated with automatic approach and avoidance tendencies toward physical activity and sedentary behaviors.

#### 4.2.1. Lateralized Readiness Potentials

LRP results showed a shorter latency of S–LRP when approaching stimuli depicting physical activity compared to sedentary behaviors, a larger positive deflection of S–LRP when avoiding stimuli depicting sedentary behaviors compared to physical activity, and a smaller positive deflection when avoiding compared to approaching stimuli depicting physical activity. These findings are consistent with the behavioral results, and showed, for the first time, that faster reaction times to approach physical activity and to avoid sedentary behaviors result from faster sensory integration (S–LRP), not faster motor planning (R–LRP) (Hypothesis 2). These results also highlight the fact that approaching physical activity and avoiding sedentary behaviors are congruent conditions (i.e., the intended response supports the required response), whereas avoiding physical activity and approaching sedentary behaviors are incongruent conditions (i.e., the intended response hampers the required response). These observations are consistent with the fact that all the participants of this study intended to be physically active and, as such, that avoiding physical activity and approaching sedentary behaviors was conflicting with their conscious goal of becoming physically active.

#### 4.2.2. Event-Related Potentials

ERP results revealed higher levels of conflict monitoring (larger early N1) and inhibition (larger N2) when avoiding stimuli depicting sedentary behaviors compared to physical activity (Hypothesis 3). These results suggest that higher levels of control were activated to counteract a general trend to approach sedentary behaviors. This finding is consistent with the proposition presented in our recent systematic review contending that behaviors minimizing energetic cost are rewarding and, as such, are automatically sought (Cheval et al., 2018). This proposition also concurs with previous work claiming that individuals possess a general trend to conserve energy and avoid unnecessary physical exertion (Lee, et al., 2016; Lieberman, 2015), thereby explaining the negative affect that could be experienced during vigorous exercise (Brand & Ekkekakis, 2018; Ekkekakis, 2017; Ekkekakis, Parfitt, & Petruzzello, 2011) and the general evaluation of physical effort as a cost (Croxson, Walton, O’Reilly, Behrens, & Rushworth, 2009; Shadmehr, Huang, & Ahmed, 2016). However, these cortical outcomes were not significantly influenced by the usual level of MVPA (Hypothesis 4). Taken together, these findings call for a cautious interpretation of the behavioral results. Faster reaction times when approaching physical activity and avoiding sedentary do not imply a general trend to approach physical activity, i.e., movement and energy expenditure, as often interpreted in the literature. Our results showed that these behavioral observations are actually associated with higher levels of inhibition likely aiming at counteracting a general trend to avoid physical exertion and enabling individuals to be more physically active.

ERP results also revealed higher levels of attentional processing (larger P1 and late N1), conflict monitoring (larger early N1), and inhibition (larger N2) when exposed to sedentary behaviors compared to physical activity stimuli, irrespective of whether these stimuli should be approached or avoided. These results are consistent with previous studies arguing that stimuli related to sedentary behaviors can represent a threatening temptation for individuals who intend to be or are physically active (like the participants of our study) as these stimuli interfere with the successful implementation of physical activity goals (Cheval, Sarrazin, Boisgontier, & Radel, 2017; Rouse, Ntoumanis, & Duda, 2013). As such, stimuli associated with sedentary behaviors may automatically trigger higher-level mechanisms preparing the individual to overcome this potential threat.

### 4.3. Strengths and Limitations

Strengths of our study include the investigation, for the first time, of the cortical activity associated with automatic approach tendencies toward physical activity and sedentary behaviors, the use of different ERP metrics that consistently showed that avoiding sedentary behaviors requires more cortical resources than avoiding physical activity, the use of LRP measures to investigate the processes occurring during sensory integration and motor preparation, the use of sophisticated EEG statistical analyses suited to examine the whole scalp throughout the duration of the response, the control of approach and avoidance tendencies toward neutral stimuli, and the validation of these results through sensitivity analyses. However, some potential limitations should also be noted. First, the usual level of physical activity was assessed using a self-reported questionnaire, which may not accurately reflect the objective level of physical activity. Yet, two independent and validated scales were used to assess physical activity and yielded consistent results. Assessing physical activity, but also sedentary behaviors, using device-based measures will be important in future research. Second, the sample size of this study was small. However, the linear mixed models used to analyze the behavioral data allowed the inclusion of all trials in the model (i.e., not the average performance per individual), which yielded an appropriate statistical power. By contrast, there was a potential power issue in the EEG analysis. In view of these two limitations (self-reported assessment of physical activity and low sample size), the non-significant effect of the level of physical activity on the cortical activity associated with automatic approach and avoidance tendencies toward physical activity and sedentary behaviors should be interpreted with caution. Third, this study involved individuals who were physically active or who intended to. Future research should examine whether the brain correlates of approach and avoidance tendencies differ between physically inactive individuals who intend and do not intend to be physically active. In the absence of intention to be active (i.e., to approach physical activity and avoid sedentary behaviors), sedentary behaviors may not be perceived as a threat. Therefore, sedentary behaviors may not affect conflict monitoring, inhibition, and motor preparation. Fourth, the neutral stimuli (i.e., square vs. circles) changed the simple effects of approaching compared to avoiding stimuli depicting physical activity and sedentary behaviors. Accordingly, interpreting these simple effects seems inappropriate. Future studies seeking to control for the automatic approach-avoidance bias toward neutral stimuli should carefully pre-test the neutral stimuli.

## 5. Conclusion

The LRP findings revealed that faster reaction times to approach physical activity and avoid sedentary behaviors were related to brain processes occurring during sensory integration, not motor preparation. The ERP findings revealed that being faster at avoiding stimuli depicting sedentary behaviors required higher levels of conflict monitoring and inhibition compared to avoiding stimuli depicting physical activity. Therefore, contrary to what behavioral results suggested, these EEG findings suggested that sedentary behaviors are attractive and that individuals intending to be active need to activate additional cortical resources to counteract this attraction.

## Acknowledgements

MPB is supported by a postdoctoral fellowship, a grant for a long-term research abroad, and research grants from the Research Foundation – Flanders (FWO; 1504015N, 1501018N). We thank Andy Katzenmeier for helping with data acquisition.

## Author Contributions

Study design: BC, RR, MPB. Data collection: BC, ET. Data analysis: BC, ET, JF, NB, DO, MPB. Draft preparation: BC, MPB. Figures preparation: JF, NB, DO, MPB. Manuscript edition: BC, ET, NB, JF, DO, RR, MPB.

## Inform Consent

Informed consent was obtained from all participants included in the study. Participants were compensated for participation.

## Ethics Approval

Ethics Committee of the University of Geneva, Switzerland approved this study.

## SUPPLEMENTARY MATERIALS

**Supplementary material 1.**
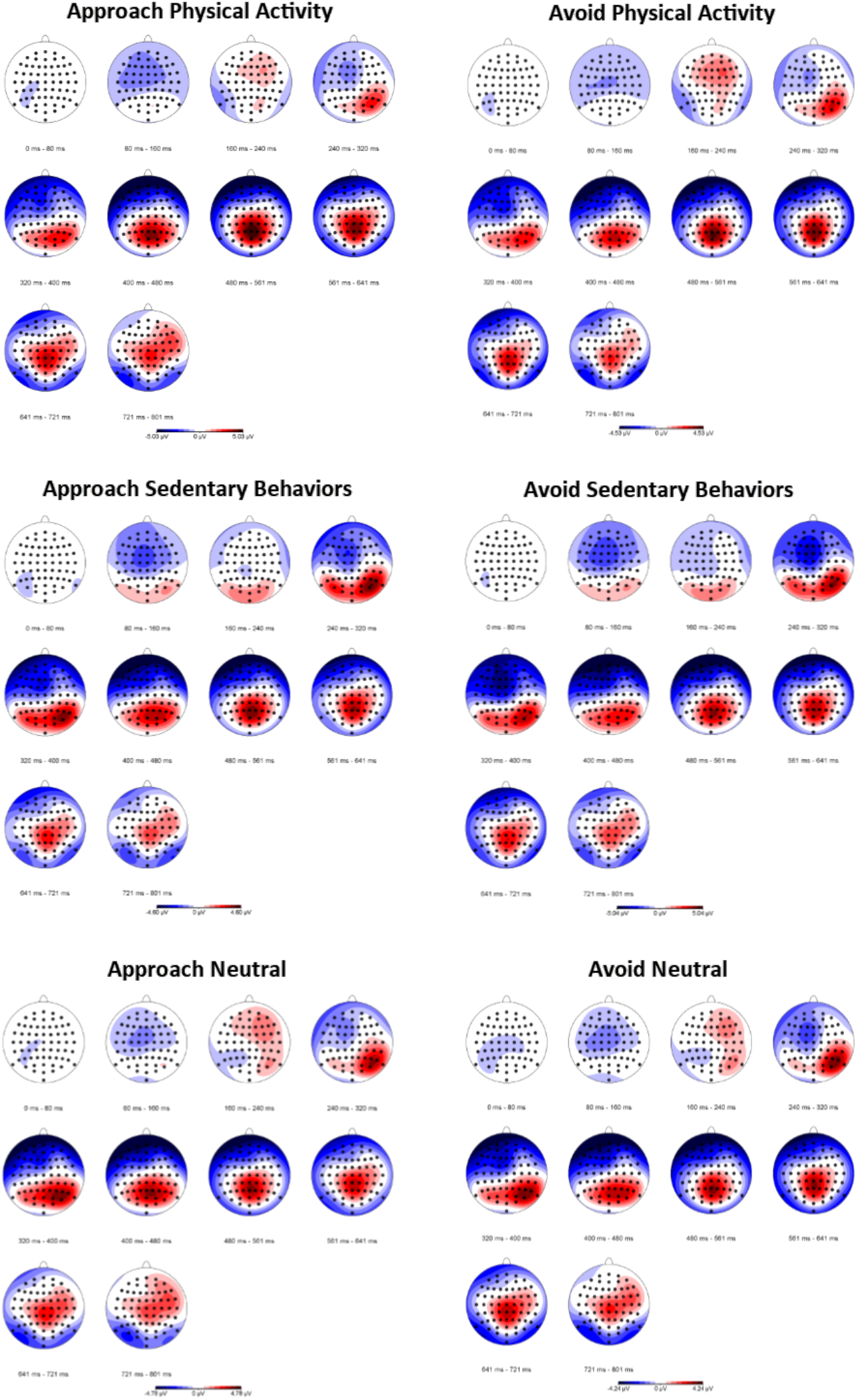
Topographical figures for all conditions.

**Supplementary material 2.**
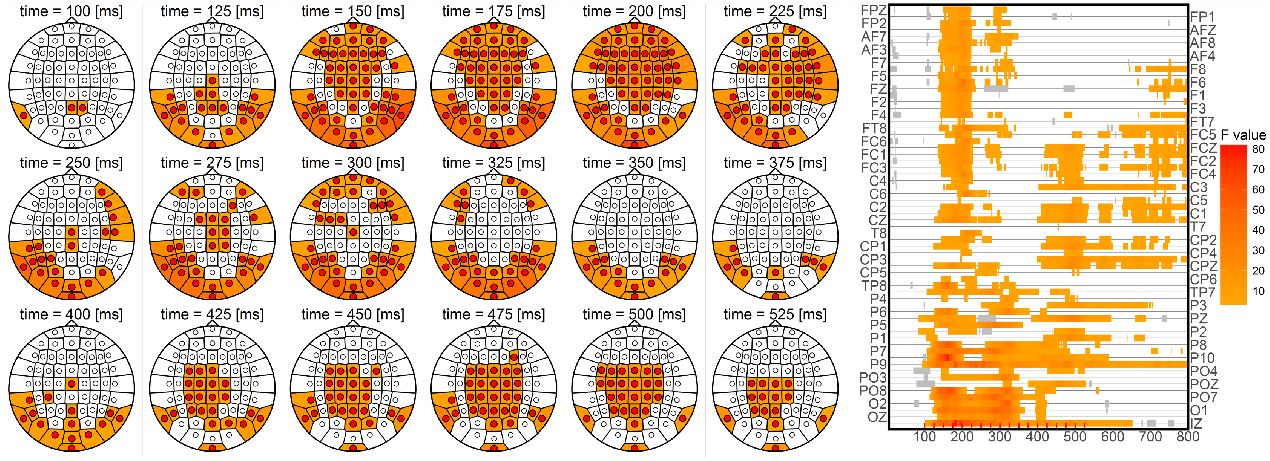
Results of the results for the simple effect of approaching stimuli depicting sedentary behaviors rather than physical activity.

**Supplementary material 3.**
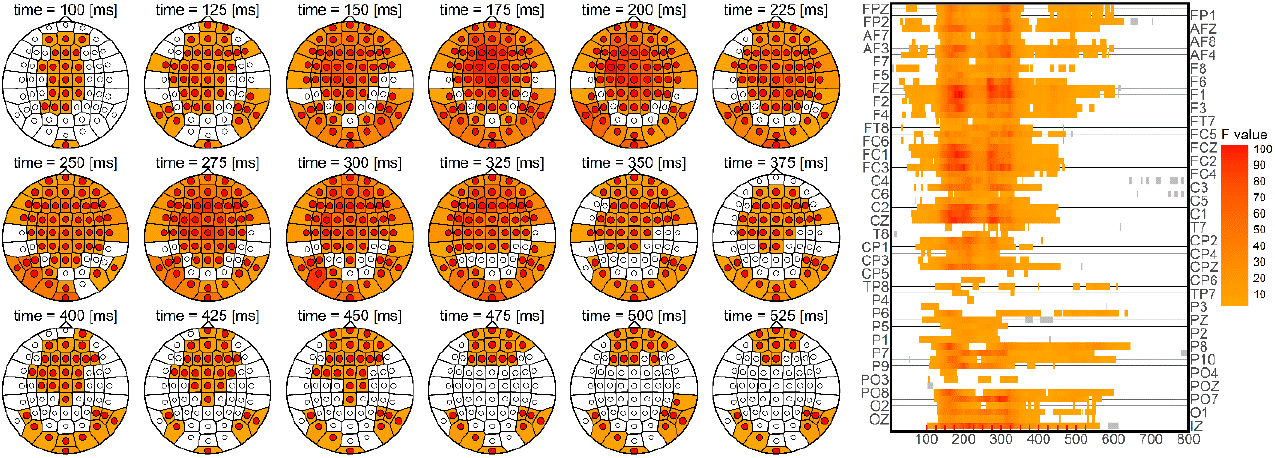
Results of the results for the simple effect of avoiding stimuli depicting sedentary behaviors rather than physical activity.

**Supplementary material 4.**
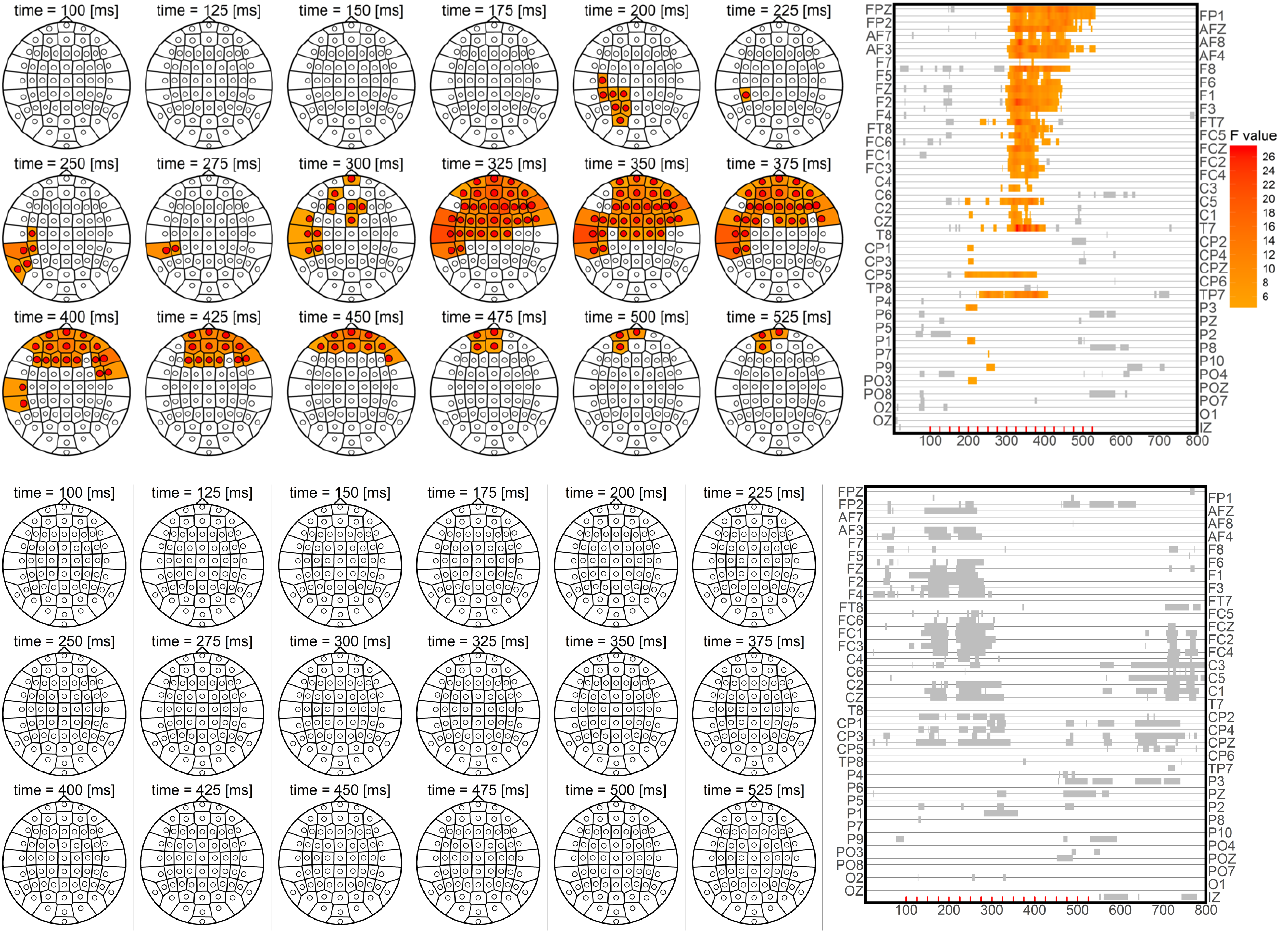
Results of the results for the simple effect of approaching rather than avoiding stimuli depicting sedentary behaviors.

**Supplementary material 5.**
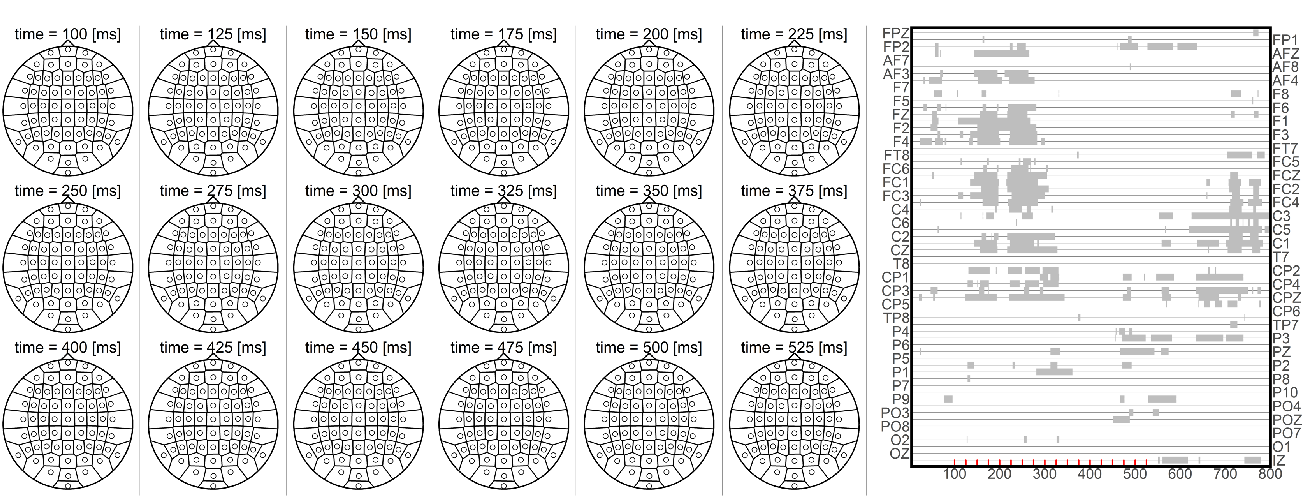
Results of the results for the simple effect of approaching rather than avoiding stimuli depicting physical activity.

**Supplementary material 6.**
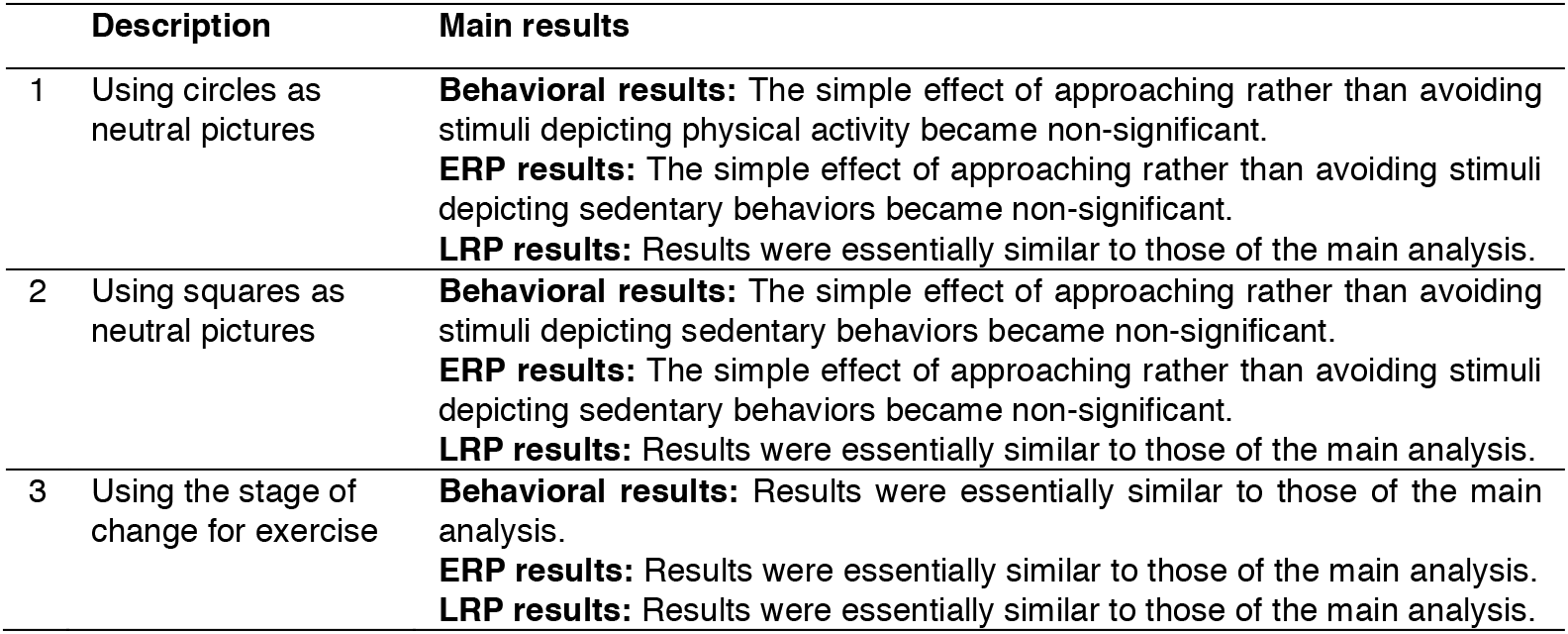
Summary of the sensitivity and complementary analyses.

